# Myc stimulates cell cycle progression through the activation of Cdk1 and phosphorylation of p27

**DOI:** 10.1101/492694

**Authors:** Lucía García-Gutiérrez, Gabriel Bretones, Ester Molina, Ignacio Arechaga, Juan C. Acosta, Rosa Blanco, Adrian Fernandez, Leticia Alonso, Piotr Sicinski, Mariano Barbacid, David Santamaría, Javier León

## Abstract

Cell cycle stimulation is a major transforming mechanism of Myc oncoprotein. This is achieved through at least three concomitant mechanisms: upregulation of cyclins and Cdks, downregulation of Cdk inhibitors p15 and p21 and the degradation of p27. The Myc-p27 antagonism has been shown to be relevant in human cancer. To be degraded, p27 must be phosphorylated at Thr-187 to be recognized by Skp2, a component of the ubiquitination complex. We previously described that Myc induces Skp2 expression. Here we show that not only Cdk2 but Cdk1 phosphorylates p27 at the Thr187, which was previously unreported. Moreover, Myc induced p27 degradation in murine fibroblasts through Cdk1 activation, which was achieved by Myc-dependent cyclin A and B induction. In the absence of Cdk2, p27 phosphorylation at Thr-187 was mainly carried out by cyclin A2-Cdk1 and cyclin B1-Cdk1. We also show that Cdk1 inhibition was sufficient for the synthetic lethal interaction with Myc. This result is relevant because Cdk1 is the only Cdk strictly required for cell cycle and the reported synthetic lethal interaction between Cdk1 and Myc.

**Summary blurb:** Myc antagonizes p27 activity in cancer. Myc activates Cdk1 to phosphorylate p27, marking p27 for degradation. This depends on Myc-mediated cyclin A and B induction. Cdk1 inhibition is sufficient for a synthetic lethal interaction with Myc

## INTRODUCTION

Progression through the cell cycle phases is under the control of a family of serine/threonine protein kinases. These kinases are heterodimers consisting of a catalytic subunit, the cyclin-dependent protein kinase (Cdk) and a regulatory subunit, the cyclin, required for the Cdk to be active. Although Cdk and cyclin are large protein families, Cdk1, 2, 4 and 6 and A, B, E, D-type cyclins are identified as the major regulators of the cell cycle [1]. Systematic knockout of *Cdk* loci in the mouse germline has shown that Cdk2 [2, 3], Cdk4 [4, 5] and Cdk6 [6] are not essential for cell cycle progression of most cell types, although loss of each of these Cdks results in particular developmental defects. Moreover, concomitant loss of the genes of interphase Cdk does not result in a general disturbance of the cell cycle in most cell types, being Cdk1 alone sufficient to drive the cell cycle [1, 7].

Myc (also called c-Myc) is an oncogenic transcription factor of the helix-loop-helix/leucine zipper protein family. Activation of transcription by Myc depends on formation of heterodimeric complexes with Max proteins. Myc-Max heterodimers bind to DNA sequences called E-boxes in the regulatory regions of its targets genes and recruit transcriptional coactivators, albeit Myc has also the ability to repress genes through less known mechanisms (for reviews see [8-10]). Myc is found deregulated in nearly half of human solid tumors and leukemia, and appears frequently associated with tumor progression [11-13].

One of the best characterized functions of the transcription factor Myc is its potent ability to induce cell proliferation, mainly by promoting the transition from G1 to S-phase stage of the cell cycle, a feature that has been linked to its pro-oncogenic activity. Indeed, the enforced Myc expression in quiescent cells is sufficient to mediate cell cycle entry. At least two major mechanisms account for this: (i) the transcriptional activation of genes whose products are required for cell cycle progression, including a number of cyclins (D2, A, E); (ii) the repression of *CDKN2B* (p15) and *CDKN1A* (p21) genes and (iii) the impairment of p27^KIP1^ (p27 here after) activity as cell cycle inhibitor (reviewed in [14]).

It is established that the Myc-p27 antagonism is one of the major mechanisms for Myc-mediated tumorigenic action. p27 is a Cdk inhibitor that is downregulated in proliferating cells and in many tumors. Cyclin E-CDK2 is considered p27’s primary target [15, 16], although other targets rather than CDK2 have been proposed [17]. The ability of Myc to overcome the p27-mediated proliferative arrest has been demonstrated in cell culture models [18, 19] and also in animal carcinogenesis models [20]. This antagonistic effect of Myc on p27 is mediated through several concomitant mechanisms: (i) Myc induces cyclin D2 and Cdk4, which sequester p27 in CDK-cyclin complexes [21, 22]; (ii) Myc induces expression of Cullin 1 (Cul1) [23] and Cks1 [24], both components of the SCF^SKP2^ complex and (iii) we showed that Skp2, the p27-recognizing subunit of a protein SCF^SKP2^ complex is a Myc target gene [25]. Moreover, Skp2 itself has been considered to have oncogenic potential and is found overexpressed in many human tumors [26, 27].

Previous studies indicated that p27 must be phosphorylated at Thr187 (human protein) to be recognized by the SCF^SKP2^ ubiquitin ligase complex, and thus to be efficiently ubiquitinated and targeted for proteasome-mediated degradation [28, 29]. In this work, we studied the mechanism of Myc-mediated phosphorylation of p27. The results unveil the pivotal role of Cdk1 on p27 phosphorylation and the relevance for Cdk1-based synthetic lethal approaches to control Myc in cancer

## RESULTS

### 1. Myc induces phosphorylation of p27 mediated by Cdk1 and Cdk2 in human leukemia cells

Previous results in our laboratory in a human myeloid leukemia cell line K562 have shown that the ability of Myc to promote cell cycle progression depends on the repression of p27 protein levels [19]. We used a K562 derivative cell line, called Kp27MER, which contains a Zn^2+^-inducible p27 construct and the chimeric protein Myc-ER that is constitutively expressed but activatable by 4-hydroxi-tamoxifen (4HT). In this model the induction of p27 arrested cell proliferation, while Myc-ER activation by 4HT induced p27 phosphorylation at the Thr-187 and partially rescued the proliferative arrest state [19, 25].

Concomitant treatment with 50 or 75 µM Zn^2+^ (to induce p27) and 200 nM 4HT (to activate Myc) for 12 hours resulted in decreased p27 levels, as expected (Fig 1A). Interestingly, MycER activation induced cyclin A2 expression, in agreement with the induction of proliferation and p27 phosphorylation (Figure 1A). The down-regulation of endogenous Myc upon 4HT addition is a marker of the MycER activation. We investigated whether Myc induced p27 phosphorylation was specific for cyclin E-Cdk2 complexes as previously reported [30, 31] or due to their redundancy, other cyclin-Cdk complexes could be involved.

**Figure 1.**
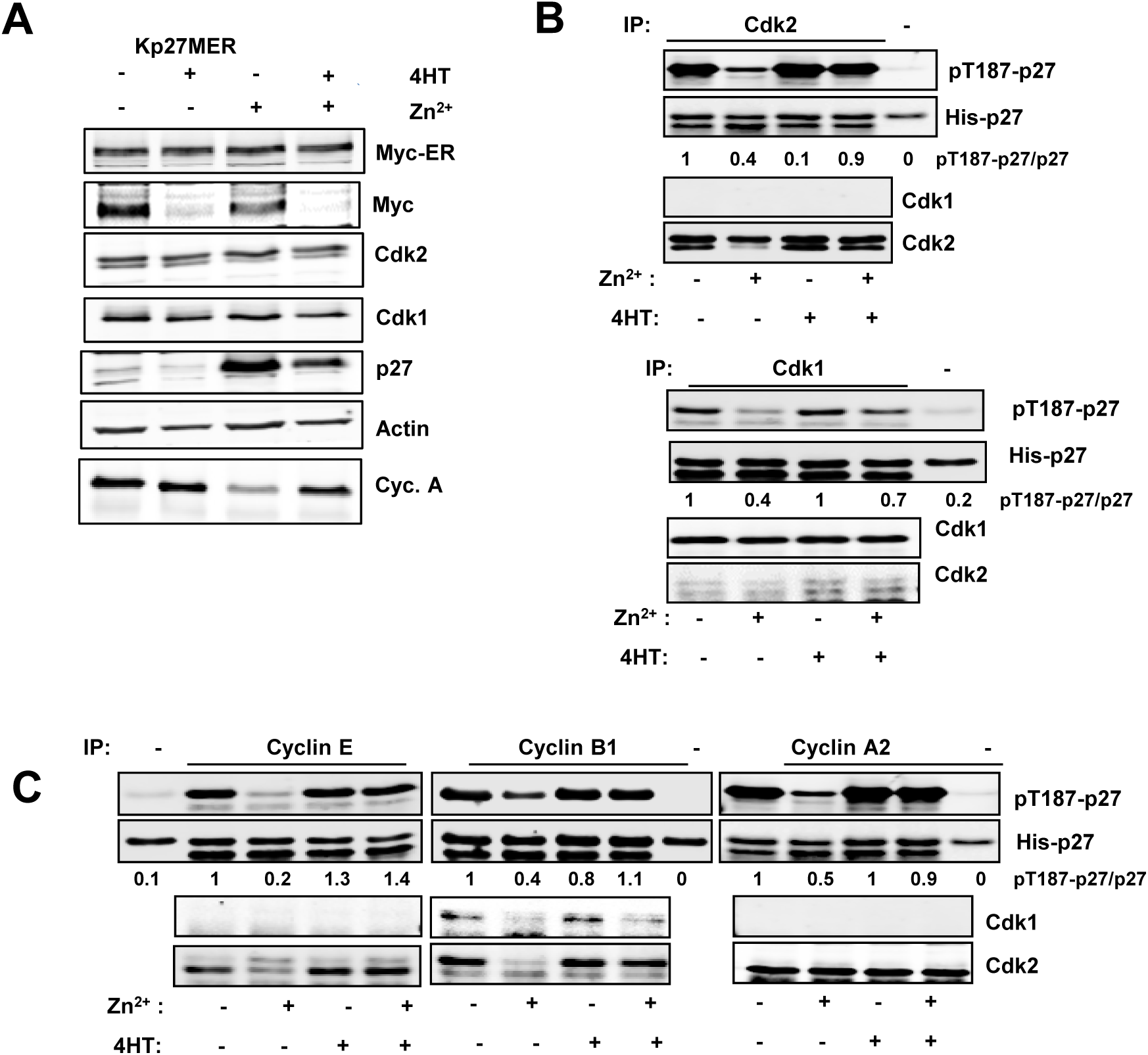
Induction by Myc of p27 phosphorylation in leukemia cells. **A)** Kp27MER cells were treated with 50 µM ZnSO4 and 200 nM 4HT for 24 hours as indicated to induce ectopic p27 expression and MycER activation. Cdk1 and Cdk2 expression and p27 levels are shown. **B)** Immunoprecipitated Cdk1 (upper panel) and Cdk2 (lower panel) complexes from Kp27MER cells treated with Zn^2+^ and 4HT for 12 h as indicated and subjected to *in vitro* kinase assays with His-p27 as substrate. Levels of Cdk1 and Cdk2 in immunoprecipitates are also shown. Relative levesl of pT187-p27 vs p27 determined by signal densitometry are shown below each lane. **C**) Immunoprecipitated cyclin B1, cyclin A2, cyclin E complexes from Kp27MER treated with Zn and 4H Zn^2+^ and 4HT for 12 yas indicated and subjected to *in vitro* kinase assays over His-p27 as substrate Levels of cyclin A2, cyclin B1, cyclin E in the immunoprecipitates are also shown.

*In vitro* kinase assays using His-p27 as substrate revealed that, after 12 hours of treatment, the activation of MycER rescued the kinase activity not only of Cdk2 (as expected) but also of Cdk1 over p27 (Fig 1B). To investigate the cyclin that activates the Cdks upon Myc activation, cyclins A2, B1 and E were immunoprecipitated from lysates of Kp27MER treated with Zn^2+^ and 4HT and subjected to *in vitro* kinase assays. The results showed that the kinase activity of the three immunocomplexes was enhanced by Myc (Fig 1C). These results suggest that not only Cdk2 but Cdk1 can phosphorylate p27 at the Thr-187 and promote its degradation.

### 2. Myc stimulates proliferation and p27 degradation in cells without cyclin E or Cdk2

We set out a series of experiments to elucidate the role of Cdk1 on Myc-dependent stimulation of cell cycle progression independently of Cdk2 activity. For this purpose, we first compared the effect of Myc enforced expression on the proliferation of wild-type MEFs, and MEFs lacking cyclin E or Cdk2 expression (wild type, *Cdk2*^-/-^ and C*cne*^-/-^ MEFs respectively). Myc enforced expression was achieved by lentiviral transduction and the elevated Myc levels in transduced cell lines was confirmed by immunoblot (Fig 2A). Comparison between the proliferation rates of *Cdk2*^-/-^cells and *Ccne*^-/-^cells with that of their overexpressing Myc counterparts showed that Myc enforced expression increases cell proliferation of *Cdk2*^*-/-*^, as well as *Ccne*^-/-^ MEFs (Fig 2B).

**Figure 2.**
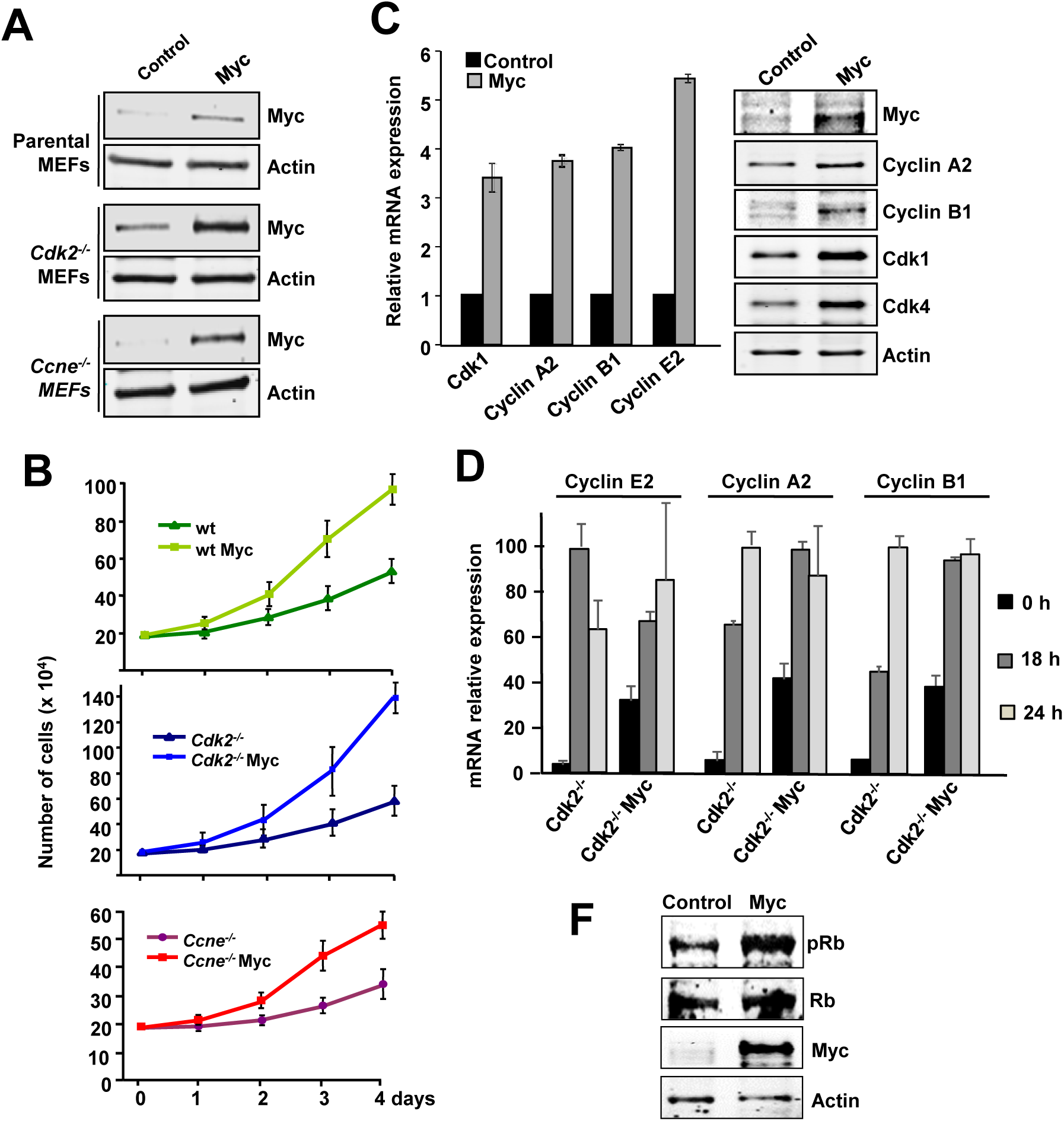
Proliferation analysis of Myc transduced *cdk2*^*-/-*^ and *ccne*^*-/-*^ MEFs. **A)** Myc transduction of parental, C*dk2*^*-/-*^ and *Ccne*^*-/-*^ MEFs analyzed by western blot. Actin protein levels were used as loading control. **B)** Proliferation curves of wild-type (wt), *Cdk2*^*-/-*^ and *Ccne*^*-/-*^ MEFs stably transduced with a constitutive Myc construct compared with controls. Data is shown as total number of cells per condition. Error bars represent s.e.m. of two independent experiments **C)** Left panel: mRNA expression of *Cdk1, Ccna2, Ccnb1* and *Ccne2* genes analyzed by RT-PCR and normalized to β actin levels. Error bars represent s.e.m. of at least two independent experiments. Right panel: Protein expression was analysed by western blot. Actin levels were used as loading control. **D**) mRNA expression of *Ccne2, Ccna2* and *Ccnb1* in serum-starved *Cdk2*^-/-^ MEFs and *Cdk2*^-/-^Myc MEFs (0 h time point) and at the indicated periods of time after serum stimulation. mRNA leves were determined by RT-PCR and made relative to the maximum expression for each cyclin. Error bars represent s.e.m. of two independent experiments. **F)** Phosphorylation of Rb protein analysed by western blot in *Cdk2*^-/-^ MEFs and *Cdk2-*/-Myc MEFs using a phospho-specific antibody for pRb.

To explore the mechanisms responsible for this Myc-mediated hyper proliferative phenotype we first studied the expression of cyclins and Cdks involved in cell cycle progression in *Cdk2*^-/-^ MEFs. The results showed that in Myc-overexpressing cells, Cdk1, Cdk4, cyclin A2, cyclin B1 and cyclin E2 expression were higher than in control cells at both, mRNA and protein levels (Fig2C). Cell starvation followed by serum stimulation of *Cdk2*^-/-^ MEFs revealed that upon Myc overexpression, the induction of cyclins at the mRNA level was accelerated in time when compared with control cells (Fig 2D). In consistency with in the increased proliferation and cyclin expression, the retinoblastoma protein was hyperphosphorylated in Myc-overexpressing cells (Fig2F). In summary, we showed that Myc overexpression induced proliferation in cells deficient in either cyclin E or Cdk2 and this was accompanied with an increase in cyclin A2 and B1 expression, as described in other models [34-37].

Due to the importance of the opposite correlation between Myc and p27 in cell cycle progression and their connection through Skp2, we analyzed the contribution of p27 (*Cdkn1B*) to Myc-induced proliferation. p27 levels were analyzed in *Cdk2*^-/-^ MEFs stably transfected with the chimeric protein Myc-ER (*Cdk2*^-/-^ MER MEFs). Cells were grown until confluence to enforce p27 accumulation and then treated with 4-hydroxi-tamoxifen (4HT) to activate the constitutively expressed Myc-ER protein. Activation of Myc-ER was accompanied by an increase in Cdk1 and Skp2, while p27 protein levels were reduced (Fig 3A). One major mechanism responsible for the Myc-induced proliferation is the degradation of p27 via proteasome. This process has been reported to rely on the phosphorylation of p27 by Cdk2 as described in the introduction. Thus, to investigate the effect of Myc on p27 stability in the absence of Cdk2, *Cdk2*^*-/-*^ Myc MEFs and their corresponding control *Cdk2*^*-/-*^ MEFs were transfected with a p27-YFP construct and treated with cycloheximide, a protein synthesis inhibitor. After different periods of cycloheximide treatment, p27 protein levels were analyzed by western blot (Fig 3B). The results showed that p27 protein levels decreased faster in cells overexpressing Myc than in control cells, suggesting that Myc affects p27 stability in the absence of Cdk2. The quantification of p27 protein levels shows that p27 stability is reduced by half in Myc overexpressing cells (Fig 3C). In summary we have shown that Myc was able to promote proliferation and p27 degradation in *Cdk2*^-/-^ MEFs.

**Figure 3.**
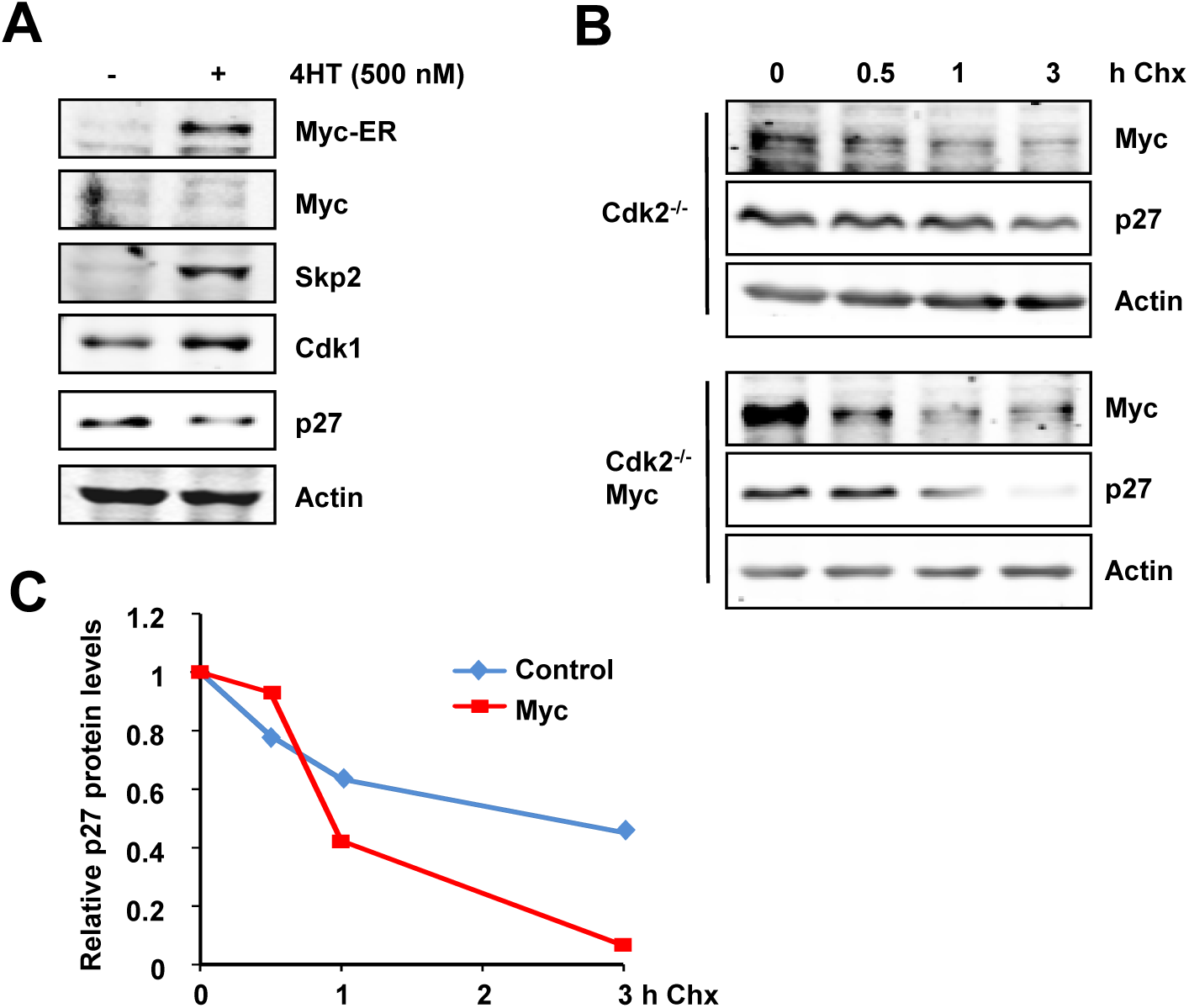
Myc-mediated degradation of p27. **A)** Protein levels of Myc, Skp2, Cdk1 and p27 of *Cdk2*^-^*/-* -MER MEFs grown until confluence and treated with 500 nM of 4HT for 18 hours (overnight). Actin levels were used as loading control. **B)** Protein stability of p27 in *Cdk2*^*-/-*^ control and *Cdk2*^*-/-*^ Myc MEFs transfected with a p27-YFP construct measured by western blot. Levels of p27-YFP were detected after 0, 0.5, 1 and 3 hours of cycloheximide treatment (30 µg/mL). β-actin levels were used as loading control. **C**) Densitometric quantification of the p27 signals of the immunoblot of panel B, normalized against β-actin levels.

Myc-mediated degradation of p27 can be explained by Myc induction of Skp2 [25] an effect that we have confirmed in MEFs (Fig 3B). However, to be recognized by Skp2, p27 must be phosphorylated at Thr-187, a phosphorylation that has been reported to be mediated by Cdk2-cyclin E [30, 31]. However, we have shown that Cdk1 could phosphorylate p27 in human leukemia cells (Fig 1B). Indeed, previous reports demonstrated that *Cdk2*^-/-^ mice showed phosphorylated p27 at the Thr187 [3]. Thus, we hypothesized that in the absence of Cdk2, Myc would induce p27 phosphorylation by activating Cdk1.

To test this hypothesis, we first explored p27 phosphorylation by Cdk1 at the Thr-187 in *Cdk2*^-/-^ MEFs. Thus, Cdk1 immunocomplexes were prepared from *Cdk2*^-/-^ Myc MEFs and *Cdk2*^-/-^ control MEFs and subjected to *in vitro* kinase assays to test their activity on p27 at the Thr-187 and whether Myc increases it. As shown in Fig 4A, Cdk1 immunocomplexes showed kinase activity over p27 and this activity was higher in *Cdk2*^-/-^Myc cells when compared to controls without Myc enforced expression. Besides, treatment of the immunocomplexes with the Cdk1 inhibitor purvalanol A lead to a decrease in p27 phosphorylation levels compared to untreated. The presence of Cdk1 in the immunoprecipitated complexes was confirmed by immunoblot (Fig 4A).

**Figure 4.**
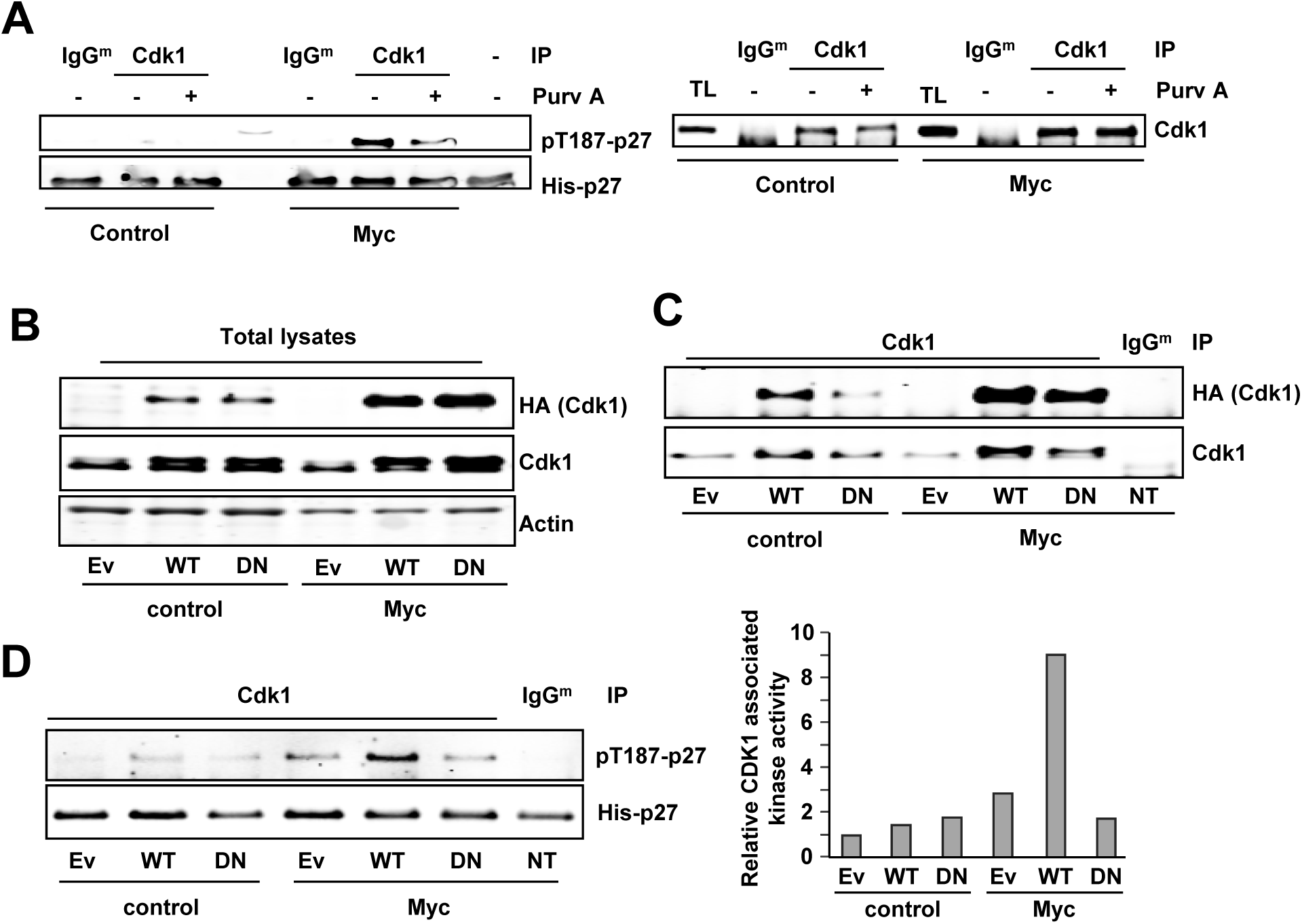
Cdk1 mediated phosphorylation of p27 at the Thr187. **A)** Left panel: *In vitro* kinase assay of Cdk1 immunoprecipitated from *cdk2*^*-/-*^ control MEFs and *cdk2*^*-/-*^ Myc MEFs using His-p27 as substrate. Phosphorylation levels of p27 at the Thr187 were detected using a phospho-specific antibody. Kinase buffer His-p27 was used as negative control and unspecific IgG as control for the specificity of the antibody used for immunoprecipitation. Immunocomplexes were treated with 10 µM purvalanol A (Purv A) or DMSO for 30 min at 30 °C prior to the kinase assay. Right panel: Levels of the immunoprecipitated Cdk1 assayed in the experiment of the left panel. **B)** Total lysates from *Cdk2*^*-/-*^ control MEFs and *cdk2*^*-/-*^ Myc MEFs transfected with empty vector (Ev) HA-Cdk1-wt (WT), HA-Cdk1-DN (DN) or non-transfected (NT). Total Cdk1 and exogenous Cdk1 are shown (anti-Cdk1 and anti-HA antibodies respectively). Actin was used as loading control. **C)** Total Cdk1 immunoprecipitated from protein extracts showed in the left panel and. Immunoprecipitated exogenous Cdk1 levels measured with anti-HA. **D)** Right panel: Kinase activity of Cdk1wild-type (WT) and Cdk1DN (DN) on Thr187 of p27. Left Panel: Densitometry quantification of the kinase assay showed in E.

These results opened the novel possibility that Cdk1 was also in charge of p27 phosphorylation and that Myc enhanced this activity. To verify that Cdk1 was able to phosphorylate p27 at the Thr-187, we transfected *Cdk2*^-/-^Myc MEFs and *Cdk2*^-/-^parental MEFs with an inactive mutant of Cdk1 lacking kinase activity (Cdk1^D146N^, Cdk1^DN^ herein after) and we performed *in vitro* kinase assays. The empty vector (EV) and the wild-type Cdk1 were used as controls. The transfected wild-type Cdk1 and Cdk1^DN^ were tagged with an HA epitope, which served to detect the exogenous Cdk1 (Fig 4B). Immunocomplexes were prepared with an anti-Cdk1 antibody, and the exogenous immunoprecipitated Cdk1 was detected by immunoblot with anti-HA antibody (Fig 4C). The kinase assays showed that overexpression of Cdk1^wt^ in *Cdk2*^-/-^ Myc MEFs led to higher phospho-p27 levels while the Cdk1^DN^ form did not show this effect when compared with the empty vector. Furthermore, overexpression of either wild-type Cdk1 or Cdk1^DN^ showed no significant differences in p27 phosphorylation compared with the empty vector in *Cdk2*^-/-^ control MEFs (Fig. 4D). This suggests that the limiting factor in p27 phosphorylation is not Cdk1 but cyclin levels, as higher Cdk1 protein levels do not have any effect unless Myc is overexpressed.

The above results demonstrated that Myc stimulates the phosphorylation of p27 mediated by Cdk1. This would be a novel function for Cdk1, as the Cdk in charge of p27 phosphorylation at Thr-187 so far reported is Cdk2 [31, 35]. Next, we investigated the cyclin partner of Cdk1 in this function. For this purpose, we used cells deficient in cyclin E1/E2. We silenced cyclin A2 expression in *Ccne*^-/-^MEFs by shRNA lentiviral transduction. Both Cdk1 and Cdk2 immunocomplexes showed reduced kinase activity on p27 *in vitro*, although Cdk1 activity was less affected by cyclin A2 downregulation than that of Cdk2 (Fig 5A). This suggests that cyclin B could be activating Cdk1 in these cells, consistently with the results showed in Figure 1. We also tested whether Myc induced p27 phosphorylation. *Ccne*^-/-^ MEFs were transduced with a Myc-IRES-GFP construct or the empty vector (Lv141) and selected by GFP expression. Cdk1 or cyclin A2 were immunoprecipitated and the *in vitro* kinase assays showed that Myc overexpression led to increased Cdk1 activity over p27 (Fig 5B), as well as increased cyclin A2-associated kinase activity (Fig 5C). Purvalanol A was used as a control of the kinases assays and in all cases abolished p27 phosphorylation. Altogether the results showed that both Cdk1 and Cdk2 were able to phosphorylate p27 at the Thr-187 in a cyclin E independent manner, suggesting that, in addition to Cdk2-cyclin A, Cdk1-cyclin A and/or Cdk1-cyclin B might be responsible for p27 phosphorylation.

**Figure 5.**
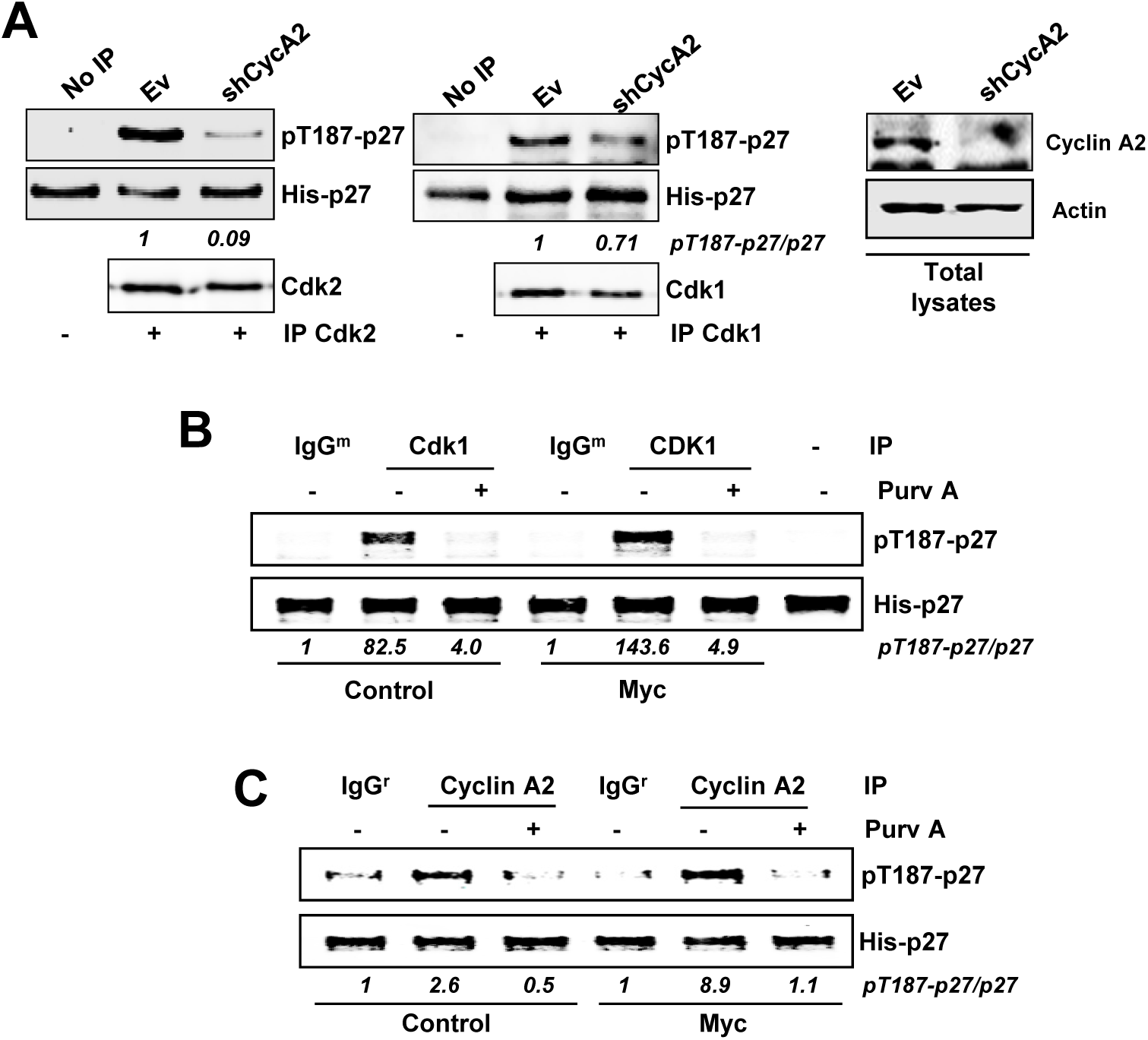
p27 is phosphorylated at the Thr187 in *Ccne*^*-/-*^ MEFs. **A)** *In vitro* kinase assays of Cdk2 and Cdk1 from *Ccne*^*-/-*^ MEFs using His-p27 as substrate. Immunoprecipitated Cdk2 and Cdk1 are shown. The silencing of cyclin A2 upon lentiviral infection is shown at the right panel. **B)** Cdk1 from *Ccne*^*-/-*^-control MEFs or *Ccne*^*-/*-^-Myc MEFs was immunoprecipitated and assayed *in vitro* over p27. Control immunoprecipitations were carried with unspecific mouse IgGs (IgG^m^). Signal densitometry quantification of the kinase assay is shown below each lane. **C)** Cyclin A from *Ccne*^*-/-*^-control MEFs or *Ccne*^*-/*-^-Myc MEFs was immunoprecipitated and assayed *in vitro* over p27. Kinase buffer with His-p27 was used as negative control and unspecific rabbit IgG (IgG^r^) as control for the specificity of the antibody used for immunoprecipitation. Immunocomplexes were treated with 10 µM purvalanol A (Purv A) or vehicle (DMSO). Control immunoprecipitations were carried with unspecific rabbit IgGs. Signal densitometry quantification of the kinase assay is shown below each lane.

Next, we tested the potential role of cyclin A as Cdk1 partner in parental and Myc-overexpressing *Cdk2*^-/-^ MEFs. Cyclin A complexes from *Cdk2*^-/-^ MEFs showed kinase activity over p27 which was higher in *Cdk2*^-/-^Myc cells. Besides, treatment of the complexes with purvalanol A decreased p27 phosphorylation (Fig 6A). Furthermore, we analyzed the requirement of cyclin A in this kinase activity. The downregulation of cyclin A2 in *Cdk2*^-/-^ and *Cdk2*^-/-^Myc MEFs was achieved by shRNA lentiviral transduction (Fig 6B). Kinase assays showed that Myc overexpression resulted in increased Cdk1 activity over p27, but cyclin A2 depletion did not reduce it (Fig 6C). The result showed that the increased kinase activity did not depend only on Myc-dependent upregulation of cyclin A2. Thus, we explored cyclin B1 as the Cdk1 activating partner to give active complexes in the phosphorylation of p27. We immunoprecipitated cyclin B1 from *Cdk2*^-/-^Myc MEFs and control *Cdk2*^-/-^ MEFs protein extracts. *In vitro* kinase assays revealed that Myc increased cyclin B1-associated kinase activity on p27 while downregulation of cyclin A expression did not affect it (Fig 6D).

**Figure 6.**
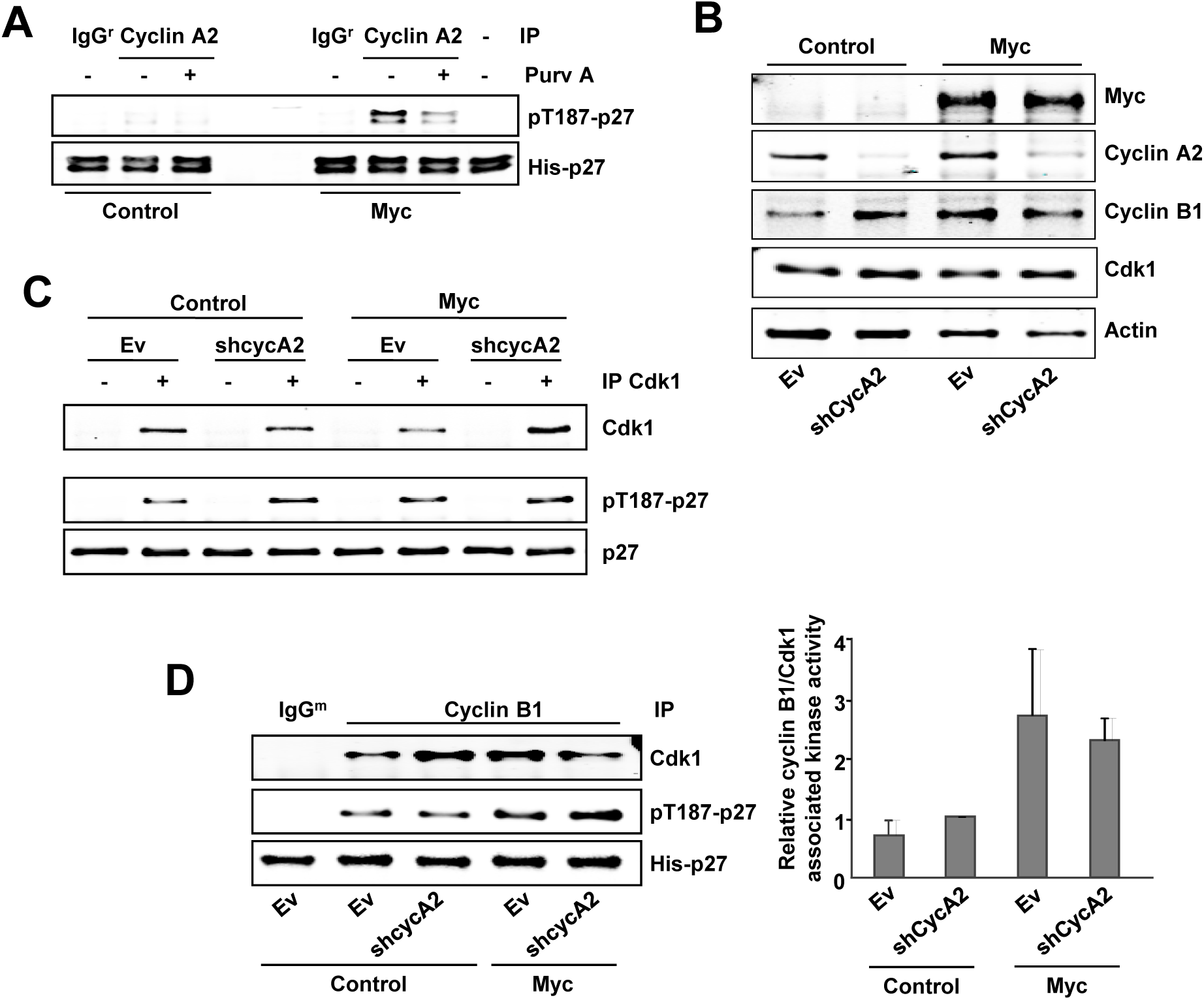
Cyclin A2 is not essential for Cdk1 to phosphorylate p27 *in vitro*. **A)** *In vitro* kinase assay of cyclin A2 complexes from *Cdk2*^*-/-*^ MEFs and *Cdk2*^*-/-*^Myc MEFs using His-p27 as substrate. Kinase buffer with His-p27 was used as negative control and control immunoprecipitation was carried out with unspecific rabbit IgG as control for the specificity of the antibody used for immunoprecipitation. Immunocomplexes were treated with 10 µM purvalanol A (Purv A) or vehicle (DMSO). **B)** Total lysates of knocked down cyclin A2 *Cdk2*^*-/-*^ and *Cdk2*^*-/-*^ Myc MEFs assayed by western blot. Cyclin A2, Cyclin B1, Cdk1 and Myc levels are shown. Actin levels were used as loading control. **C)** *In vitro* kinase assay of Cdk1 complexes from *Cdk2*^*-/-*^ and *Cdk2*^*-/-*^ Myc MEFs with knocked down cyclin A2 expression. The amount of immunoprecipitated Cdk1 is shown. **D)** Left panel: *In vitro* kinase assay of cyclin B1 complexes from *Cdk2*^*-/-*^ and *Cdk2*^*-/-*^Myc MEFs with knocked down cyclin A2 expression. Right panel: Densitometry quantification of the relative cyclin B1 kinase activity. Error bars represent s.e.m. of the quantification of two independent experiments.

### 3. Myc induces proliferation and p27 phosphorylation mediated by Cdk1

As Cdks show redundant functions within the cells, we decided to use a triple knock out MEF cell line lacking *Cdk2, Cdk4* and *Cdk6* functional genes (*Cdk2*^-/-^; *Cdk4*^-/-^; *Cdk6*^-/-^ MEFs, termed TKO MEFs herein after). This model would help to verify that Cdk1 was also responsible for the phosphorylation of p27 upon Myc expression. TKO MEFs were transduced with a Myc-IRES-GFP construct and GFP expressing cells were sorted and pooled. The resulting cell line was termed TKO-Myc MEFs.

First, comparison between TKO and TKO Myc MEFs proliferation rates we found that Myc overexpressing cells grew much faster than parental cells (Fig 7A), in agreement with an increase in the percentage of cells in S-phase (Fig 7B). Consistently, *Cdk1, Ccna2, Ccnb1* and *Ccne2* expression was elevated in cells overexpressing Myc at the mRNA (Fig 7C) and protein level (Fig 7D). Moreover, the expression of cyclins A2, B1 and E2 was not only higher in TKO-Myc than parental TKO MEFS but also induced earlier in time upon serum stimulation (Fig 7E), revealing the strength of Myc as promoter of cell proliferation, even in the only presence of Cdk1 as cell cycle Cdk. In agreement, downregulation of exogenous Myc in TKO-Myc MEFs by shRNA transduction (Fig7F) lead to arrested cell proliferation (Fig7G) and reduced expression of *Cdk1, Ccna2, Ccnb1* and *Ccne2* (Fig 7H).

**Figure 7.**
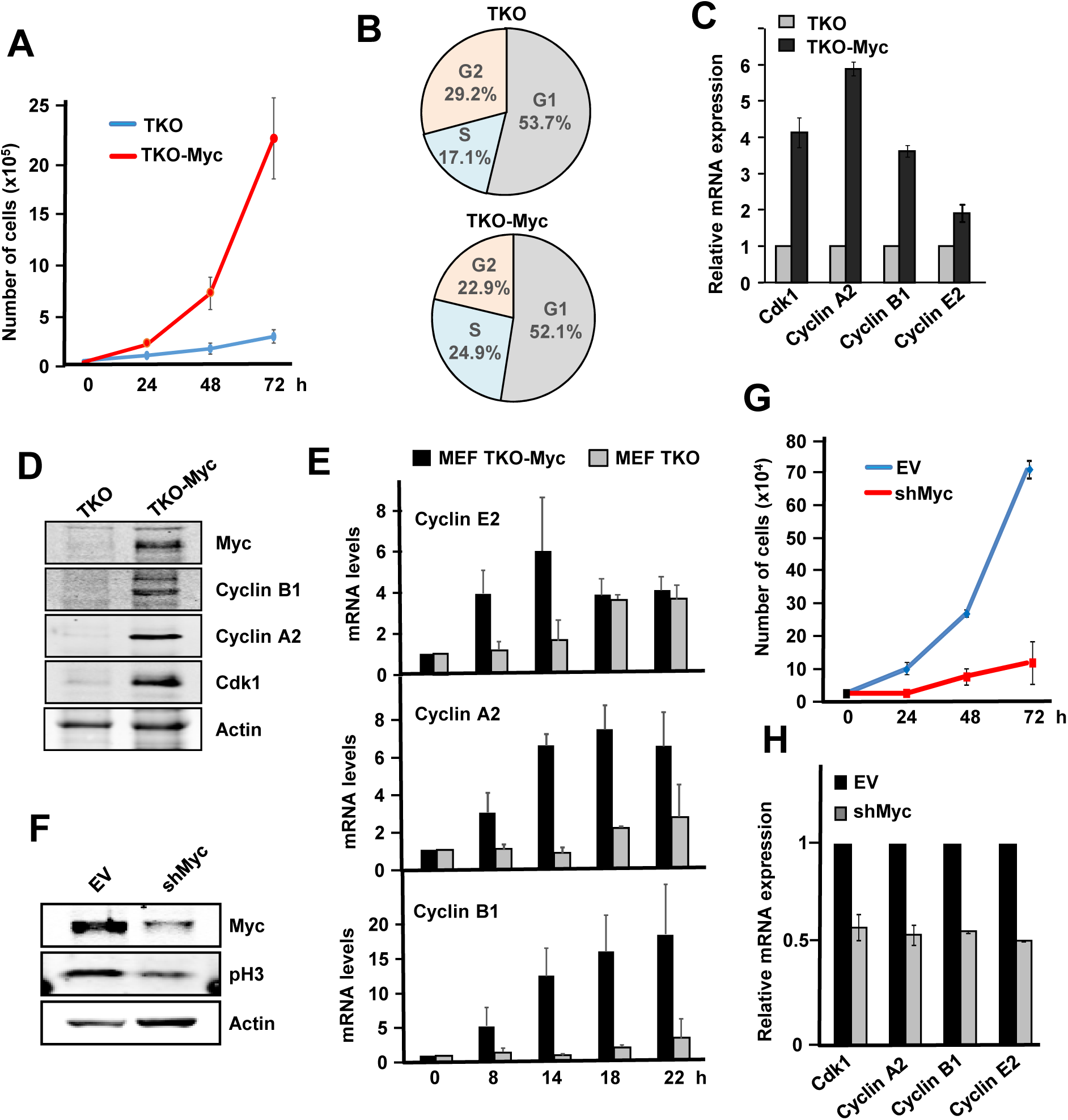
Myc induces cell proliferation and cell cycle-related genes in *cdk2*^*-/-*^; *cdk4*^*-/-*^ and *cdk6*^*-/-*^ (TKO) MEFs. **A)** Proliferation curves of TKO and TKO-Myc MEFs Error bars represent s.e.m. of three independent experiments. **B**) Cell cycle distribution in TKO and TKO-Myc MEFs. Data are mean values of two experiments with similar results. **C)** mRNA expression of C*cna2, Ccnb1, Ccne2* and *Cdk1* analyzed by RT-PCR of TKO MEFs and TKO-Myc MEFs. mRNA expression is normalized to *actin* levels. Error bars represent s.e.m. of two independent experiments. **D)** Cyclin B1, Cyclin A2, Cdk1 and Myc protein levels analysed by western blot in TKO and TKO-Myc MEFs. Actin levels are used as loading control. **E**) mRNA expression of *Ccne2, Ccna2* and *Ccnb1* at the indicated periods of time after serum stimulation in serum-starved TKO and TKO-Myc MEFs. mRNAs were determined by RT-PCR normalized to β-actin mRNA levels. Error bars represent s.e.m. of three independent experiments. **F)** TKO-Myc MEFs with downregulated Myc expression by shRNA transduction for *MYC* gene. Myc and phospho-H3 levels are shown. Actin levels are used as loading control. **G)** Proliferation curve of TKO Myc MEFs compared with Myc knocked down TKO Myc MEFs by shRNA transduction. Error bars represent s.e.m. of two independent experiments. **H**) Effect of Myc depletion in TKO cells. mRNA expression of *Ccna2, Ccnb1, Ccne2* and *Cdk1* mRNA analyzed by RT-PCR of TKO-Myc MEFs and TKO-Myc transduced with shMyc lentivirus. mRNA expression is normalized to actin levels. Error bars represent s.e.m. of two independent experiments

As the only Cdk involved in cell cycle progression of TKO MEFs is Cdk1, we wondered if Myc was able to induce Cdk1 kinase activity in these cells. We performed immunoblots using a phospho-specific antibody against the consensus amino acid sequence recognized and phosphorylated by Cdks. The results showed that Myc overexpression resulted in higher Cdk activity in TKO MEFs whereas downregulation of exogenous Myc levels in TKO-Myc MEFs led to a decrease in total Cdk activity (Fig 8A). We next used TKO MEFs to confirm that Cdk1 was able to phosphorylate p27 at the Thr187 site for further degradation. We first analyzed the expression of *Skp2*, a Myc target gene [25], needed for proteasomal p27 degradation. We observe that in TKO-Myc MEFs, *Skp2* mRNA levels were high while ectopic Myc silencing led to a decrease in Skp2 levels (Fig 8B), which is consistent with the results previously obtained using the *Cdk2*^*-/-*^ MEFs cell line. Besides, Skp2 increases and endogenous p27 decreases when Myc is overexpressed (Fig 8C). To directly test if cells lacking Cdk2, Cdk4 and Cdk6 can phosphorylate p27 at the Thr-187 through Cdk1 complexes we first prepared lysates TKO-Myc cells transfected with the p27 construct p27YFP and the results showed phosphorylated p27 (Fig 8D). As a control, TKO-Myc MEFs transfected with p27YFP were treated with roscovitine, a Cdk2/Cdk1 inhibitor. After 6 hours of roscovitine treatment, pT187-p27 levels were highly reduced compared with that of untreated cells (Fig 8D). Altogether, the results are consistent with the hypothesis that the phosphorylation is carried out by Cdk1. To corroborate this, Cdk1 immunocomplexes prepared from TKO and TKO-Myc MEFs were subjected to *in vitro* kinase using His-p27 as substrate. The results showed that overexpression of Myc (TKO-Myc cells) induced Cdk1 kinase activity on p27 (Fig 8E). To determine which is the cyclin that activates Cdk1 in TKO-Myc cells we performed immunoprecipitations with for cyclin A2, cyclin B1 and cyclin E as well as Cdk1 and determined the kinase activity on p27. Immunocomplexes of both cyclin A2 and cyclin B1 contained Cdk1 in TKO MEFs and were able to phosphorylate p27 *in vitro*. Immunoprecipitated cyclin E from TKO-Myc MEFs did not contain Cdk1 and thus, showed no kinase activity (Fig 9A). In order to determine which of the two complexes (Cdk1-cyclin A2 or Cdk1-cyclinB1) was more active over p27. For this, we carried out kinase assay with three different amounts of the immunoprecipitated complexes (Fig 9A). The kinase assays demonstrated that cyclin A2 and cyclin B1 immunocomplexes showed similar p27 phosphorylation levels *in vitro*, although the amount of Cdk1 that co-immunoprecipitated with cyclin A2 is lower than with cyclin B1. This result suggested that cyclin A2-Cdk1 complexes were more efficient in phosphorylating p27 at the Thr-187 than cyclin B1-Cdk1 complexes. However, complexes obtained by immunoprecipitating Cdk1 yielded much lower p27 phosphorylation levels compared to the previous ones, although Cdk1 amounts were much higher.

**Figure 8.**
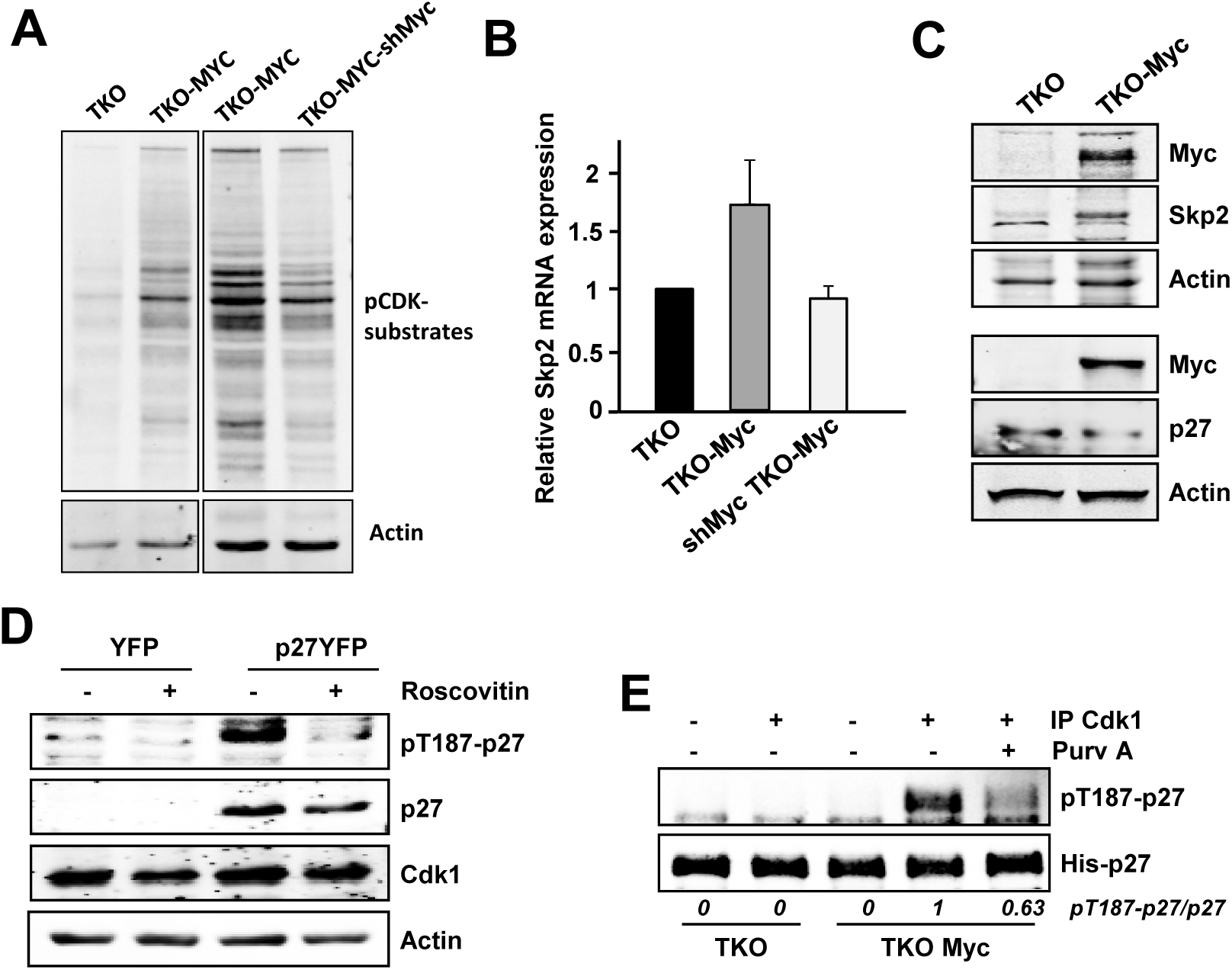
Myc induces p27 downregulation and phosphorylation in TKO MEFs. **A)** Protein extracts from TKO, TKO Myc and shMyc TKO Myc MEFs showing total Cdk activity measured by western blot using a phospho-specific antibody for the consensus sequence recognized and phosphorylated by Cdks. Actin levels are used as loading control. **B)** mRNA expression of *Skp2* in TKO, TKO-Myc and shMyc-TKO-Myc MEFs. mRNA expression is normalized to β *actin* levels. Error bars represent s.e.m. of two independent experiments. **C)** Protein levels of Myc, p27 and Skp2 from TKO comparing to TKO-Myc MEFs are shown. Actin levels were used as loading control. **D)** pT187-p27, total p27, Myc and Cdk1 levels analysed in protein extracts from YFP-p27 transfected TKO-Myc MEFs and treated with 10 µM roscovitine for 6 h. Actin levels were used as loading control. **E)** *In vitro* kinase assays of Cdk1 immunocomplexes from TKO and TKO-Myc MEFs on His-p27 as substrate. Immunocomplexes were treated with 10 µM purvalanol A (Purv A) or the corresponding amount of vehicle (DMSO) prior to the kinase assay. Relative levels of pT187-p27 vs p27 determined by signal densitometry are shown below each lane.

**Figure 9.**
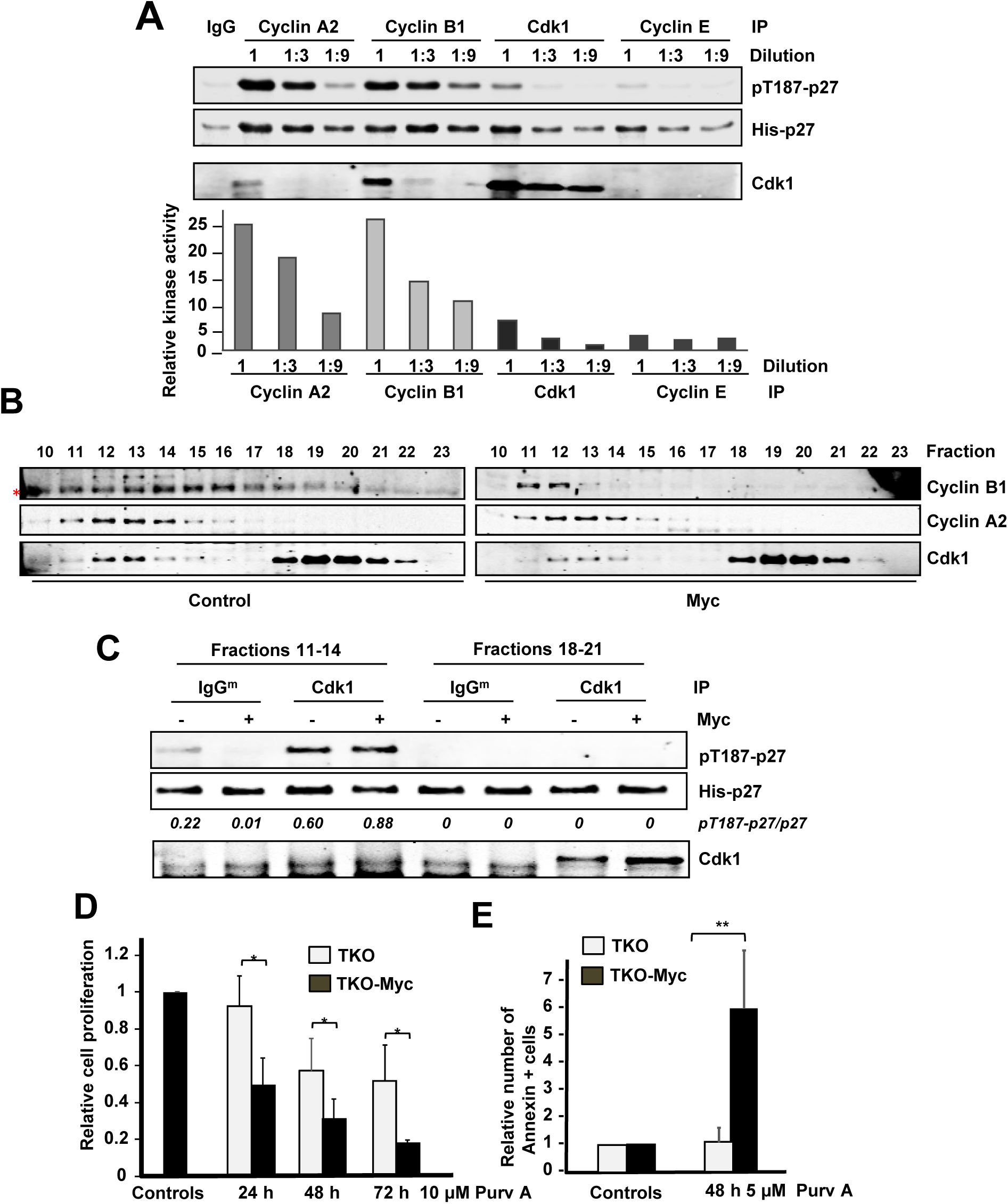
Cyclin A2 and cyclin B1 associated to Cdk1 phosphorylate p27 at the Thr187 in MEFs lacking Cdk2. **A)** *In vitro* kinase assays of cyclin A2, cyclin B1, cyclin E2 and Cdk1 complexes from TKO-Myc MEFs using His-p27 as substrate. The immunoprecipitated complexes were undiluted or diluted in buffer 1:3 and 1:9 before the kinase assay as indicated at the top. Quantification of the relative kinase activity of each complex is represented. **B)** Protein extracts from *Cdk2*^*-/-*^-control and *Cdk2*^*-/-*^-MYC MEFs were separated by gel-filtration chromatography and distribution of cyclin A2, cyclin B1, Skp2 and Cdk1 was analyzed by western blot (* indicates an unspecific band). **C)** Cdk1 immunoprecipitation from fractions corresponding to free Cdk1 (18-21) and complexed cyclin A2/B1-Cdk1 (11-14) *in vitro* kinase assays on His-p27 as substrate. Relative levels of pT187-p27 vs p27 determined by signal densitometry are shown below each lane. **D**) Myc increases the proliferation arrest induced by purvalanol A in TKO cells. Purvalanol A was added at 10 µM to TKO and TKO-Myc MEFs, and the proliferation was assessed by crystal violet staining after 24, 48 and 72 h of treatment. Error bars represent s.e.m. of three independent experiments. (E) Myc increases the fractio of apoptotic cells induced by purvalanol A. Cells were treated for 24 h with 5 μM purvalanol A and annexin V binding assayed Error bars represent s.e.m. of three independent experiments

To explain these results, we hypothesized that only a small fraction of the immunoprecipitated Cdk1 was forming complexes with cyclin A2 or cyclin B1, while the rest was unbound and thus, inactive Cdk1. To test this hypothesis, total protein lysates from C*dk2*^*-/-*^Myc MEFs and their control *Cdk2*^*-/-*^MEFs were separated by gel filtration chromatography. The fractions containing Cdk1 were identified by immunoblot and the results showed that Cdk1 is distributed in high-molecular mass complexes corresponding to a molecular mass of about 80 kDa (Fig 9B, fractions 11 to 14), and free form corresponding to a molecular mass of about 35kDa, i.e., the molecular mass of Cdk1 (Fig 9B, fractions 18 to 21). The cyclins B1 and A2 coeluted with Cdk1 in the high molecular mass fractions, which likely contain the cyclin-Cdk complexes, but the amount of Cdk1 in the cyclin fraction is much lower than the free Cdk1 (Fig 9B). We hypothesized that most of the Cdk1 immunoprecipitated was inactive because it was not bound to any cyclin. The fractions containing the free form of Cdk1 and the ones containing the high-molecular mass complexes of Cdk1, respectively, were pooled and Cdk1 was immunoprecipitated. The in vitro kinase assays of the immunoprecipitates over p27 showed Cdk1 in the free form was inactive while the Cdk1 from the high-molecular mass complexes showed kinase activity over p27 in vitro (Fig 9C). These results revealed that most of the Cdk1 was unbound to cyclins, but as expected, only the bound Cdk1 showed kinase activity on p27. Based on these data we conclude that Myc can trigger p27 phosphorylation through the induction of Cdk1 activity independently of Cdk2 and Cdk4/6. It has been reported the synthetic lethality previously shown for Cdk1 and Myc based on the selectivity of purvalanol against Cdk1 [38, 39]. We wanted to test this a Cdk2-, Cdk4- and Cdk6-deficient cells, comparing the effect of purvalanol A on TKO and TKO-Myc cells. In these cells the drug will only inhibit Cdk1. The results showed that purvalanol inhibited more efficiently the growth of cells overexpressing Myc (Fig 9D). TKO-Myc cells exposed to showed a dramatic increase in annexin V binding, indicating that Myc expressing cells are prone to cell death in response to Cdk1 inhibition (Fig 9D). The results suggest the lethal synthesis between Cdk1 and Myc in this model.

### 4. Conditionally *cdk1* knock out cells show that p27 phosphorylation is mediated by Cdk1-cyclin B1 and induced by Myc

As described in the Introduction, Cdk1 has been reported as the only indispensable Cdk in animal cells, i.e., no other interphase Cdk is capable of supply Cdk1 to accomplish an entire cell division [7]. To confirm the Cdk1-mediated phosphorylation of p27 at the Thr-187, we used a conditional knock out MEF cell line (*Cdk1*^*lox/lox*^ MEFs) and its respective control cell line (*Cdk1*^*+/lox*^ MEFs). Treatment of the *Cdk1*^*lox/*lox^ cell line with 4HT leads to *Cdk1* knock out, whereas the C*dk1*^*+/lox*^ cell line shows normal Cdk1 expression [40]. We first performed kinetics and found that after 72 hours of tamoxifen treatment, Cdk1 protein levels were almost undetectable, while control cells or cells treated with the vehicle (DMSO) did not show any change in Cdk1 levels (Fig 10A). No significant changes in Cdk2, cyclin A2 and B1 expression were found in Cdk1-depleted cells (Fig 10A)

Next, the effect of Myc overexpression on Cdk1 kinase activity was analyzed. For this purpose, cells were transiently transduced with a Myc-IRES-GFP construct or the corresponding empty vector. Myc overexpression after 72 hours was confirmed by immunoblot and was accompanied with increased expression of cyclin A2 and cyclin B1 (Fig 10B). We assayed the kinase activity on p27 of cyclin B1 immunoprecipitates from *cdk1*^lox/lox^ MEFs and *Cdk1*^+/lox^ MEFs. In this immunoprecipitates, the kinase activity should be due to Cdk1-cyclin B1 dimers. p27 phosphorylation was determined after 72 hours of 4HTtreatment (or vehicle) and Myc-IRES-GFP transduction. Cyclin B1 complexes were able to phosphorylate p27 *in vitro*, in agreement with our previous results and its activity was increased by Myc overexpression (Fig 10C). Cdk1 depletion resulted in totally abolition of cyclin B1-associated kinase activity and Myc was not able to overcome it (Fig 10C). This result demonstrates that cyclin B1-associated kinase activity over p27 is exclusively mediated through Cdk1, discarding the possibility that other Cdk could substitute for Cdk1 loss. When Myc is overexpressed, Cyclin B1 complexes obtained from *cdk1*^*+/lox*^ MEFs yielded higher pT187-p27 levels while 4HT treatment did not affect it (Fig 10C). We also tested the effect of Myc overexpression on the kinase activity associated to cyclin A2 after Cdk1 ablation. The results showed that Myc overexpression led to higher cyclin A2-associated kinase activity over p27 in control (DMSO) cells while knocking out Cdk1 led to higher cyclin A2 kinase activity, which was enhanced by Myc (Fig 10D). The overexpression of Myc three days after lentiviral transduction and ablation of Cdk1 in the inputs are assessed by western blot (Fig 10E)

**Figure 10.**
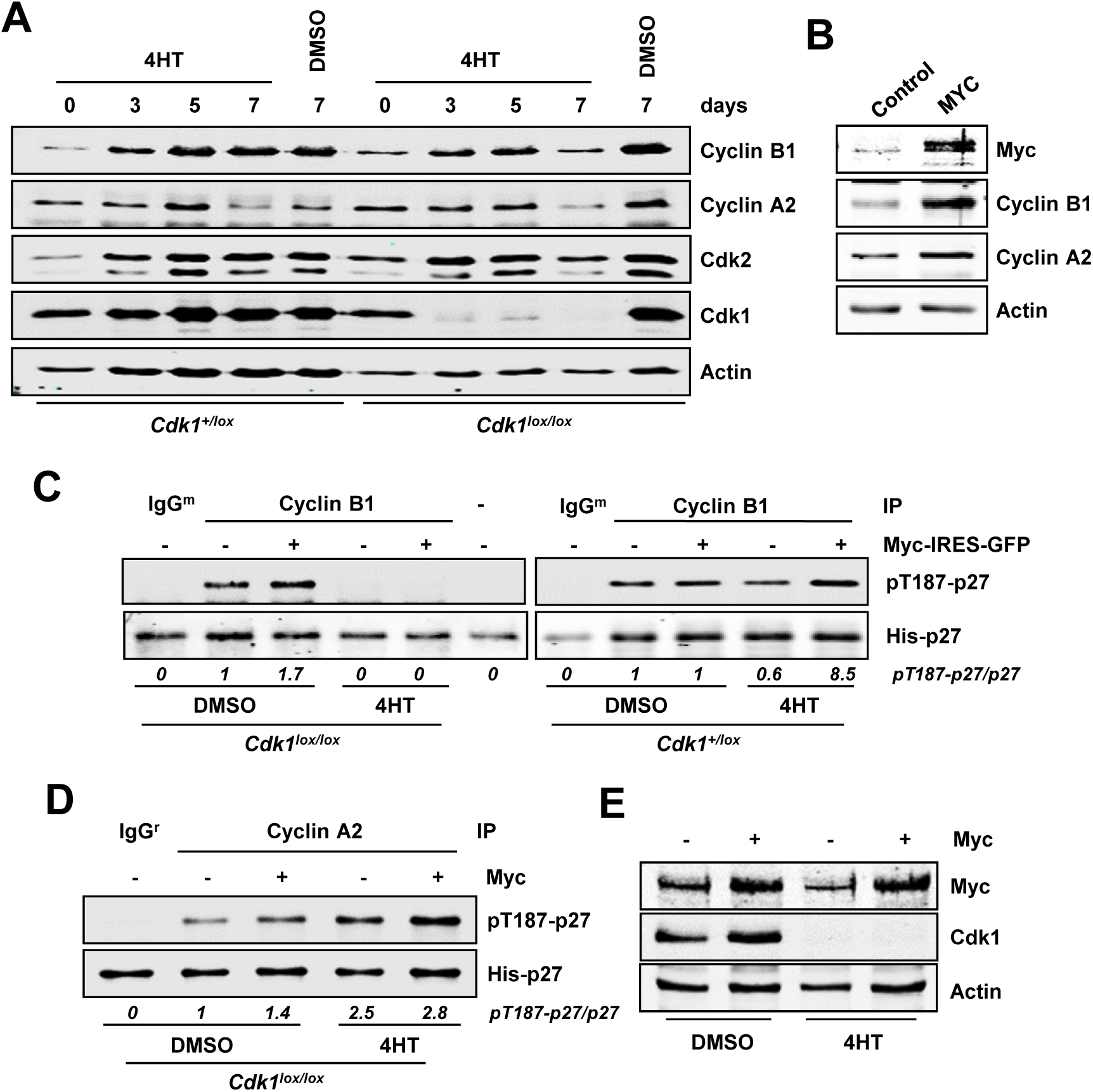
Conditional knock out of *cdk1* totally abolishes cyclin B-dependent phosphorylation of p27. **A)** Protein extracts from *Cdk1* conditional knock out MEFs and controls treated with 0.6 μM 4HT or vehicle (DMSO) for different periods of time. Expression of cyclin A2, cyclin B1, Cdk1 and Cdk2 are shown. Actin levels were measured as loading control. **B)** *Cdk1*^*lox/lox*^ MEFs transduced with a MYC-IRES-GFP construct or the corresponding control. Myc overexpression assayed by western blot. **C)** *In vitro* kinase assay of cyclin B1 immunocomplexes on His-p27 in *Cdk1*^*lox/lox*^ and control *Cdk1*^*+/lox*^ of 4HT for 72 hours (lentiviruses and 4HT were added at the same time) using p27 as substrate. Relative levels of pT187-p27 vs p27 determined by signal densitometry are shown below each lane. **D)** *In vitro* kinase assay of immunoprecipitated cyclin A from *Cdk1*^*lox/lox*^ MEFs transduced with a Myc-IRES-GFP construct or the corresponding control and treated with 4HT for 72 hours. Signal densitometric quantification of the kinase assay is shown below each lane. **E)** Expression of Myc and Cdk1 in *Cdk1*^*lox/lox*^ MEFs transduced with a Myc-IRES-GFP construct and treated with 0.6 μM 4HT or DMSO for 72 hours (lentiviruses and 4HT were added at the same time). Actin levels were determined as loading control.

## DISCUSSION

Taken together, the results shown in this work indicate that Myc induces the phosphorylation of p27 at Thr187 through the activation of not only Cdk2 (as previously described) but also through Cdk1. Cyclin B1 has Cdk1 as the unique partner capable of phosphorylating p27 in agreement with the fact that Cdk1 is being reported to show functions that cannot be compensated by any other [7]. Furthermore, kinase activity on p27 associated to cyclin A2 increased after Myc overexpression. In this case, p27 phosphorylation would be carried out by the mixture of Cdk1 and Cdk2 activities, as they both are cyclin A2 partners. In summary the results with the cells with conditional Cdk1 ablation confirmed the Myc-Cdk1-p27 pathway. However, ablation of Cdk1 lead to higher cyclin A2 activity on p27. In conditions of Cdk1 knock out, Cdk2 do not compete with other kinase for binding cyclin A2 and it leads to higher cyclin A2 -associated kinase activity. This suggests that Cdk2 is more efficient in phosphorylating p27 than Cdk1.

Our data unveils a novel mechanism by which Myc induces p27 degradation. It has been previously shown that *Skp2* is a Myc target gene and that, in order to be recognized by the ubiquitin ligase E3 complex SCF^SKP2^, p27 must be phosphorylated in Thr187. This phosphorylation was previously reported to be carried out *in vitro* by the heterocomplex cyclin E-Cdk2 [30, 35]. However, in the present work we have confirmed that it can also be mediated by Cdk1.

Which is the cyclin activating Cdk1 in Myc-expressing cells? We show here that Myc can induce expression of cyclins A, B and E but only cyclin A2 and cyclin B1 would activate Cdk1 to phosphorylate p27 in the absence of Cdk2. Taken together, our results MEFs suggest that both cyclin A2 and cyclin B1 are contributing to p27 phosphorylation through Cdk1. This is consistent with previous reports describing that cyclin B1-Cdk1 can phosphorylate p27 *in vitro* [29, 41]. Nevertheless, when we compared the activity of cyclin A2 or cyclin B1 immunocomplexes with Cdk1 immunocomplexes, we found that those with Cdk1 were much less efficient in phosphorylating p27. This can be explained if most of the Cdk1 is unbound to cyclins, and thus inactive, and this is indeed the case. Finally, the Myc-Cdk1-p27 axis is not only operative in mouse fibroblasts but also confirmed in human myeloid leukemia cells with conditional expression of Myc and p27.

In summary, Myc would promote p27 degradation through a triple effect: (i) the induction of Skp2 expression, which is the major pathway for p27 ubiquitination and degradation; (ii) the activation of Cdk2 via the upregulation of its cyclin partners and (iii) the activation of Cdk1, a mechanism previously unreported. These mechanisms are summarized in Fig 11A. In tumor cells the high levels of Myc, or the induction of Myc levels by mitogenic stimuli in normal cells, would led to p27 phosphorylation, ubiquitination and degradation. This pathway along the cell cycle is schematized in Fig 11B. Synthetic lethality has been proposed as an alternative antitumoral therapy when the oncogene involved is not easily druggable. This would be the case of Myc [42]. It has been previously identified the chemical inhibition of Cdk1 and Cdk2 to be synthetic-lethal with Myc or N-Myc overexpression [38, 39, 43]. As it is well known the apoptotic effect of Myc when cells are exposed to suboptimal growth conditions [44] thus it is conceivable that inhibition of Cdk1 would result in elevated p27 and thus an enhanced apoptotic effect of Myc. In conclusion, the results described here are relevant because, first, the opposite correlation between Myc and p27 levels is found in many human tumors and is related to poor outcome [45, 46], and second, because it provides a mechanism for the described synthetic lethality between Myc overexpression and Cdk1 pharmacological inhibition.

**Figure 11.**
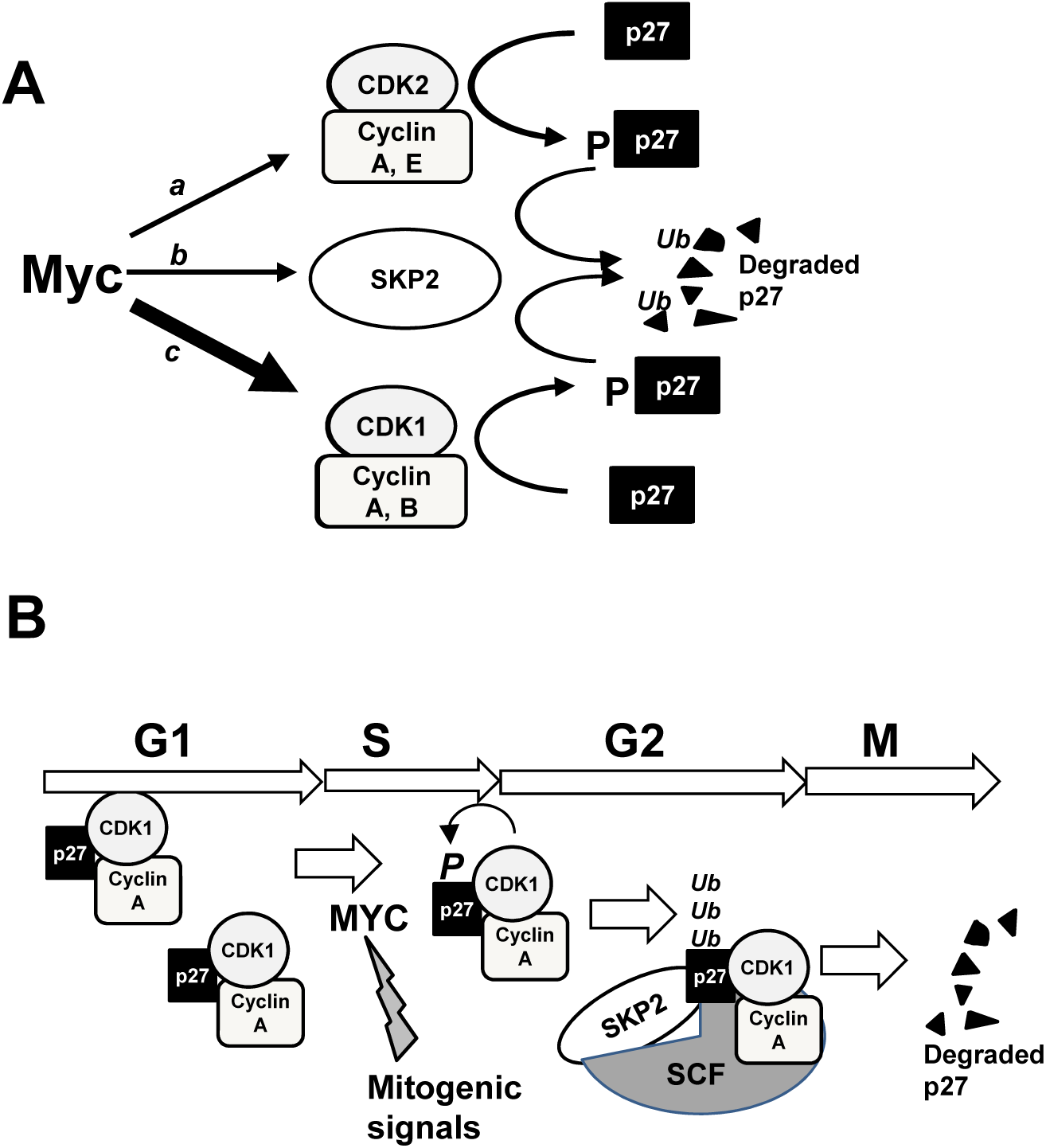
**A)** Schematic model of the Myc pathways leading to p27 degradation. These are (a) the induction of Cdk2 activity; (b) the induction of Skp2 and (c) and the induction of Cdk1 activity, shown in this work. **B)** Summary of p27 complexes and interactions and degradation along the cell cycle

## MATERIALS AND METHODS

### Cell culture, proliferation assays and transfections

Mouse embryonic fibroblast (MEFs) cell lines and Kp27MER cells were grown in DMEM (Lonza) or RPMI (Lonza) respectively, supplemented with 10% (v/v) fetal bovine serum (Lonza), 150 μg/ml gentamycin and 2 μg/ml ciprofloxacin. The following MEFs were transduced with the lentiviral vector Lv141-GFP-Myc (Genecopoeia) or its respective Lv141-GFP empty vector to generate stably or transiently Myc-overexpressing cells: *Cdk2*^*-/-*^ MEFs (derived from *cdk2*^*-/-*^ mice-G418); TKO MEFs (derived from *cdk2*^*-/-*^,*cdk4*^*-/-*^,*cdk6*^*-/-*^ mice-G418/puromycin/Hygromycin); *Ccne*^*-/-*^ MEFs (derive from *ccne1/2*^*-/-*^ mice); *Cdk1*^*lox/lox*^ MEFs (derived from *cdk1*^*lox/lox*^ mice). The following stably transfected or transduced cells were maintained in presence of the appropriate antibiotics: *Cdk2*^*-/-*^MER MEFs were grown with puromycin 3 µg/mL; pLKO-*Cdk2*^*-/-*^ MEFs and shCyclin A2 *Cdk2*^*-/-*^ MEFs were grown with puromycin 6 µg/mL; Kp27MER [19] cells were grown with puromycin (0.5 µg/mL) and G418 (250 mg/mL). Cells were grown in normoxia in a humidified atmosphere at 37°C and 5% CO_2_ except for TKO MEFs that were grown under hypoxia (3-4 % O_2_). Proliferation assays were performed as follows: cells were seeded in six-well plates at 2.5 × 10^4^ total number of cells per well (same number of wells as counting days). Cells were harvested and counted in NucleoCounter^®^ NC-100™ system (Chemometec). Sample 0 hours corresponds to 48 hours after lentiviral infection. Drug treatment concentrations: 0.6 µM for MEFs and 0.2 µM for Kp27MER of 4-hydroxi-tamoxifen (Sigma-Aldrich), 30 µg/mL of cycloheximide (Sigma-Aldrich), 10 µM of roscovitine (Sigma-Aldrich), 50 µM of Zn_2_SO_4_ and 10 µM purvalanol A. Transfection of MEFs was performed using PEI reagent (Polysciences, Inc). Briefly, PEI and DNA were mixed in free serum DMEM in a ratio 2.5:1 PEI:DNA (µg), vortexed and incubated 10-30 min RT. Culture media was replaced by half the amount of serum-free media and mixture of PEI+DNA was added to cells at 60-70% of confluency. After 12 hours of transfection mixture of PEI+DNA was removed and cells were supplemented with complete media. In order to synchronize the cells for cell cycle studies, the cells were seeded at a 60% confluence, washed with PBS and starved for two days with DMEM without FBS. After two days, the media were removed, fresh media was added with 50 µM hydroxyurea (Sigma) and further incubated for 12 hours. Then, the cells were washed with PBS and stimulated with DMEM supplemented with 10% FBS. After different incubation periods of time, cells were harvested for RNA preparation and cell cycle assay. For cell cycle assay, one million of cells were resuspended with PBS, 3mM EDTA and fixed with cold absolute ethanol overnight. Then, Hoechst stain (5 µg/mL) was added and the cells were incubated for 30 min at 37°C. Cell cycle profile was determined in MACSQuant VYB (Miltenyi Biotec) and the results were analyzed with Flow Logic software (Miltenyi Biotec).

### Lentivirus production and cell transduction

HEK293T cells were transfected with PEI as previously described to generate lentiviral particles. Mixture of packaging plasmids (pCMV-VSV-G and psPAX2 from Addgene) and construct of interest was performed in a ratio of 1:3:4 (VSV-G:psPAX2: lentiviral construct). Lentiviral particles-containing supernatants were collected and stored at 4°C in two rounds (48 and 72 hours after transfection). Supernatants were clarified at 1500 rpm for 10 minutes and filtered. For lentivirus concentration, clarified supernatants were mixed with PEG8000 (final concentration of 15% PEG8000), homogenized by inversion and led at 4°C for 24-72 h The mixture was centrifuged at 1500*xg* for 30 minutes at 4°C, and the pellet with lentivirus was dissolved in small volumes of serum-free media and stored at -80°C. For lentivirus tittering, HeLa cells were infected with increasing amounts of concentrated lentiviral particles (usually from 0.1 to 10 µL) and selected with puromycin. A multiplicity of infection (MOI) ≥ 5 was used to transduce MEFs. For infection, the cells were pelleted and resuspended in the corresponding volume of lentivirus with 8 µg/mL of polybrene. The mixture of cells and lentivirus was incubated at 37°C for 1 hour, resuspended every 10 minutes and plated in half of the volume for the corresponding plate in serum-containing media. After 12-18 hours, the same volume of media was added to reach the final volume of the corresponding plate and 48 hours after transduction lentivirus-containing media was replaced by fresh media and puromycin selection when indicated.

### Annexin V binding assay

Cells were seeded in a 60 % confluence and 12 h later treated with 5 µM Purvalanol A for 24 h. Cells were harvested, washed twice with PBS-3 mM EDTA, resuspended in 100 µL of Binding Buffer (10 mM Hepes/NaOH, 140 mM NaCl and 2.5 mM CaCl_2_, pH 7.4) and 2 µL FITC Annexin V (BD Bioscience) and incubated for 30 min at 4°C. Cells were then washed twice with PBS and resuspended in 250 µL of PBS. Annexin V binding was assayed in MACSQuant VYB (Miltenyi Biotec) and the results were analyzed with Flow Logic software (Miltenyi Biotec).

### mRNA expression analysis by qPCR

RNA extraction was carried out using Tri Reagent (Sigma-Aldrich). Roughly, 5 × 10^5^ cells were lysed in 0.5 mL of Tri Reagent^®^, 0.1 mL of chloroform was added and gently mixed. The mixture was incubated RT for 2 minutes and centrifuged at 13000 rpm for 15 min. Nucleic acid containing aqueous phase was transferred to a new 1.5 mL tube. For RNA precipitation, 0.25 mL of isopropanol was added, mixed by inversion, incubated at RT for 10 minutes and centrifuged for 15 minutes at 13,000 rpm 4°C. Supernatant was discarded and 0.5 mL of 70% ethanol added. The RNA containing pellet was mixed by vortexing and centrifuged for 5 minutes at 7500 rpm 4°C. Supernatant was discarded, RNA air dried and resuspended in RNAse free water. RNA concentration was determined by using a NanoDrop and 200 ng were resolved in an agarose gel to check the integrity of the RNA. For cDNA conversion, the iScript™ cDNA Synthesis Kit from Bio-Rad was used following manufacturer’s protocol for 1µg of RNA as template. SYBR^®^ Select Master Mix from Applied Biosystems™ was used to amplify cDNA in a CFX Connect™ Real-Time PCR Detection System from Bio-Rad. qPCRs were analyzed with the CFX Manager™ software. The mRNA expression of genes of interest was normalized to β-actin mRNA expression. Murine primer sequences used (forward and reverse primers in 5’→3’ direction) *Cdk1*: CGGCGAGTTCTTCACAGAG and AACCGGAGTGGAGTAACGAG; *Cdk4*: GGCCTGTGTCTATGGTCTGG and TTCAGCCACGGGTTCATATC; *Ccna2*: GCCAGCTGAGCTTAAAGAAAC and AACGTTCACTGGCTTGTCTTC; C*cnb1*: GACGTAGACGCAGATGATGG and GCCAGTCAATGAGGATAGCTC; *Ccne2*: GCATTCTGACCTGGAACCAC and GGAAGCAATGAACAATGAGG; *Skp2*: AATCTGCACCCAGACGTGAC and TGGAGCACTCGGACAGAATC; *bactin*: AGACTTCGAGCAGGAGATGG and AGTTTCATGGATGCCACAGG.

### Protein levels analysis by western blot

Adherent cells were directly lysed in the plate after washing once with cold PBS, while suspension cells were pelleted, washed once with cold PBS and the cell pellet lysed in the tube. Roughly, 100 µl of 1% NP40 lysis buffer (50 mM Tris-HCl pH 8, 150 mM NaCl, 1 mM EDTA, 10 mM NaF, 1% NP40 (v/v), 0.1% SDS; protease (Calbiochem) and phosphatase (Sigma-Aldrich) inhibitors added immediately before use) were used for 5 × 10^5^ cells. All the steps were performed at 4°C. Adherent material was washed once with cold PBS and directly lysed in the plate using a scrapper and protein extracts were collected in a 1.5 mL tube. Suspension cells were pelleted, washed once with cold PBS and incubated in lysis buffer for 30 min on ice, mixing them every 10 min by pipetting. Protein samples were sonicated (10 cycles in a Bioruptor^®^ Plus Sonicator device) and finally clarify by centrifugation at 14000 rpm for 20 min at 4°C. The supernatant was transferred to a new tube and kept frozen until used. Protein quantification was carried out using the Qubit^®^ Protein Assay Kit in a Qubit ^®^ 2.0 Fluorometer. Samples were resolved by SDS-PAGE and transferred to a nitrocellulose membrane. Blocking was carried out at room temperature using 4% BSA in TBS-T for 1 hour. Primary antibodies were used diluted in TBS-T 1%BSA at a final concentration of 1:1000 unless indicated. IRDye800/tolRDye680 secondary antibodies were used and signals were recorded with and Odyssey^®^ Infrared Imaging Scanner (LiCor^®^ Biosciences). Quantification and densitometry analysis was carried out using the ImageJ software. Primary antibodies: cdc2 p34-(17) (Santa Cruz Biotechnology, sc-54, monoclonal antibody used for immunoprecipitations and immunoblots); cdc2 p34 C19 polyclonal (sc-954, used for immunoblots); cdk2 M2 (sc-163); cdk4 C22 (sc-260); cyclin A2 H432 (sc-239); cyclin B1 GNS1 (sc-245, used 1:500); cyclin E H111 (sc-248, used 1:500); His-probe H15 (sc-813); myc N262 (sc-764, used 1:2000 5% BSA); p27 C19 (sc-528); p27 C19 (sc-528-G); skp2 p45 H435 (sc-7164); actin I19 (sc-1616). Phospho-Cdk substrate motif (Cell Signalling, 9477), Phospho-Thr187 p27 (Invitrogen 71-7700; used 1:500 in western blots).

### Immunoprecipitation and ***in vitro* kinase assays**

Cell lysis was performed using of non-denaturing lysis buffer (50 mM Tri-HCl pH7.5, 150 mM NaCl, 1 mM EDTA, 0.5 % NP40; protease (Calbiochem) and phosphatase (Sigma-Aldrich) inhibitors added immediately before use). After cell lysis and protein quantification as previously indicated, 50 µg of protein extracts were separated for whole cell extract analysis and 500 µg were used for each immunoprecipitation. Dynabeads^®^-protein G (Invitrogen^®^) were used to capture protein-antibody immunocomplexes. Briefly, Dynabeads (15 µL per IP) were washed twice with washing buffer (50 mM Tri-HCl pH7.5, 150 mM NaCl, 1 mM EDTA, 0.5 % NP40), added to the protein extract plus antibody mix and incubated overnight at 4°C with rotation. Next, Dynabeads^®^-protein G-immunocomplexes were collected using a magnet DynaMag™ (Invitrogen), washed four times with washing buffer and resuspended in 30 µL of 2x SDS-PAGE loading buffer. Protein-protein interactions were analyzed by western blot as previously described. Normal IgG was used as negative control for each immunoprecipitation reaction. Alternatively, 1/3 of Dynabeads^®^-protein G-immunocomplexes were used for *in vitro* kinase assay. They were additionally washed twice with kinase buffer (50 mM Hepes-NaOH pH 7.2, 150 mM NaCl, 10 mM MgCl2, 2.5 mM EGTA, 1 mM EDTA, 1 mM DTT, 10 % glycerol, 10 mM β-glycerophosphate and 10 mM NaF), resuspended in 30 µL kinase buffer supplemented with 50 µM ATP and 1 µg of recombinant His6-p27 (His-p27) protein and incubated for 30 min at 30 °C. The kinase reaction was stopped by adding 5xSDS-PAGE loading buffer to the reaction. Samples were boiled and phosphorylation of p27 at the Thr-187 was measured by western blot using a phospho-specific antibody anti phospho-Thr187 p27 (Invitrogen 71-7700). When indicated, the immunocomplexes were incubated with 10 µM Purvalanol A (or DMSO as control) in the presence of ATP for 30 min at 30 °C. Afterwards, the His-p27 was added to each reaction, mixed, incubated for 30 min at 30 °C and analyzed as described. Normal IgG and kinase buffer supplemented with ATP and substrate were used as negative controls for the kinase reaction. The relative kinase activity was determined by densitometric quantification of the western blot using the ImageJ software and represented as the fraction of pT187-p27 signal relative to total p27.

### Gel-filtration chromatography of total protein extracts

10^7^ cells were lysed as described in 200-300 µL non-denaturing chromatography lysis buffer (50 mM Hepes-NaOH pH 7.2, 150 mM NaCl, 0.05 % Triton X-100, 1 mM EDTA, 2.5 EGTA, 1 mM DTT, 10 % Glycerol and 1 mM PMSF; protease (Calbiochem) and phosphatase (Sigma-Aldrich) inhibitors added immediately before use), incubated on ice for 30 min and protein extracts clarified at 14000 rpm for 20 min t 4°C and transferred to a new tube. 200 µL of protein extracts were directly applied onto a Superdex 200 10/300 GLcolumn (GE Healthcare) pre-equilibrated with chromatography buffer (50 mM Hepes-NaOH pH 7.2, 150 mM NaCl, 0.05 % Triton X-100, 1 mM EDTA, 2.5 Mm EGTA, 1 mM DTT, 10 % Glycerol and 1 mM PMSF) and subjected to fast-performance liquid chromatography in an Äkta purifier apparatus (GE Healthcare) with a flow rate of 0.4 mL/min at 4°C. The protein complexes eluting from the column with different volume retentions according to their molecular size were fractionated in 500 µL fractions. 40 µL of each collected fraction were mixed with 10 µL of 5X SDS-PAGE loading buffer and analyzed by western blot as indicated. Alternatively, the selected fractions containing the protein complexes were mixed and subjected to immunoprecipitation and *in vitro* kinase assay as previously described.

## Acknowledgements

The work was supported by grant SAF2017-88026-R from MINECO, Spanish Government, to JL and MDD (partially funded by FEDER program from European Union). LGG was recipients of F.P.I. fellowship from Spanish Government. We are grateful Sandra Zunzunegui for technical assistance and John Sedivy and M Dolores Delgado for helpful discussions

## Author contributions

L García-Gutiérrez: formal analysis, investigation and writing-review and editing

G Bretones: formal analysis and investigation

E Molina: investigation

I Arechaga: investigation and methodology

JC Acosta: investigación and methodology

R Blanco: data curation and investigation

A Fernandez: investigation

L Alonso: investigation

P Sicinski: resources,

M Barbacid: conceptualization, formal analysis

D Santamaría: conceptualization, formal analysis

J León: conceptualization, formal analysis, funding acquisition and writing

